# Restoration of susceptibility to amikacin by 8-hydroxyquinoline analogs complexed to zinc

**DOI:** 10.1101/598912

**Authors:** Jesus Magallón, Kevin Chiem, Tung Tran, María S. Ramirez, Verónica Jimenez, Marcelo E. Tolmasky

## Abstract

Gram-negative pathogens resistant to amikacin and other aminoglycosides of clinical relevance usually harbor the 6’-*N*-acetyltransferase type Ib [AAC(6’)-Ib], an enzyme that catalyzes inactivation of the antibiotic by acetylation using acetyl-CoA as donor substrate. Inhibition of the acetylating reaction could be a way to induce phenotypic conversion to susceptibility in these bacteria. We have previously observed that Zn^+2^ acts as an inhibitor of the enzymatic acetylation of aminoglycosides by AAC(6’)-Ib, and in complex with ionophores it effectively reduced the levels of resistance *in cellulo*. We compared the activity of 8-hydroxyquinoline, three halogenated derivatives, and 5-[N-Methyl-N-Propargylaminomethyl]-8-Hydroxyquinoline in complex with Zn^+2^ to inhibit growth of amikacin-resistant *Acinetobacter baumannii* in the presence of the antibiotic. Two of the compounds, clioquinol (5-chloro-7-iodo-8-hydroxyquinoline) and 5,7-diiodo-8-hydroxyquinoline, showed robust inhibition of growth of the two *A. baumannii* clinical isolates that produce AAC(6’)-Ib. However, none of the combinations had any activity on another amikacin-resistant *A. baumannii* strain that possesses a different, still unknown mechanism of resistance. Time-kill assays showed that the combination of clioquinol or 5,7-diiodo-8-hydroxyquinoline with Zn^+2^ and amikacin was bactericidal. Addition of 8-hydroxyquinoline, clioquinol, or 5,7-diiodo-8-hydroxyquinoline, alone or in combination with Zn^+2^, and amikacin to HEK293 cells did not result in significant toxicity. These results indicate that ionophores in complex with Zn^+2^ could be developed into potent adjuvants to be used in combination with aminoglycosides to treat Gram-negative pathogens in which resistance is mediated by AAC(6’)-Ib and most probably other related aminoglycoside modifying enzymes.

## Introduction

Among many mechanisms bacteria have evolved to resist antibiotics, enzymatic modification is one of the most efficient [1, 2]. In the case of aminoglycosides, bactericidal antibiotics used to treat a wide range of bacterial infections, the most relevant mechanisms of resistance in the clinics are enzymatic inactivation by acetylation, nucleotidylation, or phosphorylation [2-4]. Although more than hundred aminoglycoside modifying enzymes have been identified in bacterial pathogens, the acetyltransferase AAC(6’)-Ib, which mediates resistance to amikacin and other aminoglycosides, is the most widespread among Gram-negative clinical isolates [5-7]. The progressive acquisition of this gene is eroding the usefulness of amikacin as well as other aminoglycosides. One way to overcome this problem is the design of new antimicrobials such as the recent introduction of plazomicin [8]. However, since this is a slow and expensive process and resistance will inevitably develop against the new antibiotics, these efforts must be complemented by other strategies to prolong the useful life of existing drugs [2, 3, 9-11]. In the case of aminoglycosides, in addition to design of new molecules [8, 12, 13], there is active research to find inhibitors of expression of aminoglycoside modifying enzymes [14-18] and to design enzymatic inhibitors [2, 3, 10, 11, 19-22]. A recent breakthrough in the search for inhibitors of enzymatic inactivation of aminoglycoside was the finding that Zn^+2^ and other metal ions inhibit the acetylation of aminoglycosides mediated by AAC(6’)-Ib *in vitro* [23]. Although concentrations beyond toxic levels were needed to interfere with resistance in growing bacteria, further research showed that the action of the metal was enhanced when complexed to ionophores, in which case low concentrations were sufficient to overcome resistance in several aminoglycoside-resistant bacteria [23-26]. We recently showed that two classes of ionophores, clioquinol (5-chloro-7-iodo-8-hydroxyquinoline)(CI8HQ) and pyrithione (N-hydroxypyridine-2-thione), when complexed to Zn^+2^ or Cu^+2^, significantly reduce the levels of resistance to amikacin in *Escherichia coli, Klebsiella pneumoniae*, and *Acinetobacter baumannii* strains harboring the *aac(6’)-Ib* gene [24-26]. CI8HQ and other substituted 8-hydroxyquinolines are being tested as treatment for cancer, neurodegenerative conditions such as Alzheimer’s, Parkinson’s, and Huntington’s diseases, and lead poisoning [27-30]. The ongoing studies and uses of these compounds indicate that human toxicity is not a serious impediment in their development as drugs for diverse diseases [29, 31]. These facts make CI8HQ and other substituted 8-hydroxyquinolines excellent candidates to be used in combination with aminoglycosides in the treatment of resistant infections. In this work we compared the effect of commercially available substituted 8-hydroxyquinolines complexed to Zn^+2^ on growth of amikacin-resistant *A. baumannii* clinical isolates.

## Materials and methods

### Bacterial strains and reagents

The *A. baumannii* A155, A144, and Ab33405 clinical isolates were used in growth and time-killing experiments to test the ability of the ionophores complexed to zinc to reduce resistance to amikacin. All three strains are resistant to amikacin but only A144 and A155 naturally carry *aac*(*6*′)-*Ib* [32-34]. Ionophores and amikacin sulfate were purchased from MilliporeSigma. [Acetyl-1-^14^C]-Acetyl Coenzyme A was purchased from Perkin-Elmer.

### Enzymatic acetylation assays

Acetylation activity was assessed using the phosphocellulose paper binding assay as described previously [35, 36]. Amikacin and [Acetyl-1-^14^C]-Acetyl Coenzyme A were used as substrates in reactions carried out in the presence of the soluble content of cells that were disrupted by sonication as described previously [37]. The reactions were carried out in a final volume of 25 μl containing 200 mM Tris-HCl, pH 7.6, 200 μM amikacin, 0.5 μCi [Acetyl-1-^14^C]-Acetyl Coenzyme A (specific activity, 60 mCi/mmol), and the enzymatic extract (120 μg protein). The reaction mixtures were incubated at 37°C for 1 h and then 20 μl were spotted on phosphocellulose paper strips. The unreacted radioactive donor substrate was eliminated from the phosphocellulose paper by submersion in 1 l hot water (80°C) followed by several washes with water at room temperature. The phosphocellulose paper strips were allowed to dry before determining the radioactivity.

### Growth inhibition and time-kill assays

The inhibition of growth of *A. baumannii* strains by amikacin and ionophore-zinc complexes was tested inoculating 100-μl Mueller-Hinton broth in microtiter plates with the specified additions using the BioTek Synergy 5 microplate reader [23]. The cultures were carried out at 37°C with shaking and contained dimethyl sulfoxide (DMSO) at a final concentration of 0.5%. The optical density at 600 nm (OD_600_) of the cultures was determined every 20 minutes for 20 h. Time-kill assays were carried out as described before [38]. Briefly, cells were cultured to 10^6^ cfu/ml in Mueller-Hinton broth. At this point the indicated concentrations of amikacin, ionophore, and zinc were added, and the cultures were continued at 37°C with shaking. Samples were removed at 0, 4, 8, 20, and 32 h, serially diluted, plated on Mueller-Hinton agar, and incubated at 37°C for 20 hours to determine the number of cfu/ml.

### Cytotoxicity assays

Levels of cytotoxicity were determined using the LIVE/DEAD Viability/Cytotoxicity Kit for mammalian cells (Molecular Probes) as described [39]. HEK 293 cells plated at a density of 10^3^ cells/well were cultured overnight under standard conditions in flat bottom, 96-well, black microtiter plates. The compounds being tested, dissolved in dimethyl sulfoxide (DMSO), were then added to the cells at increasing concentrations as indicated, and incubation was continued. As control DMSO was added to duplicate wells at same final concentration reached when adding the compounds being tested. After 24 h, the cells were washed with sterile D-PBS and incubated with the LIVE/DEAD reagent (2 μM ethidium homodimer 1 and 1 μM calcein-AM) for 30 min at 37 °C, and the fluorescence level at 645 nm (dead cells) and 530 nm (live cells) was measured. The percentage of dead cells was calculated relative to the cells treated with DMSO. The maximum toxicity control was determined using cells incubated in the presence of 0.1% Triton X-100 for 10 min. Experiments were conducted in triplicate. The results were expressed as mean ± SD of three independent experiments.

## Results

Combination therapies consisting of an antibiotic and an inhibitor of resistance can be an invaluable tool in the search for solutions to the multidrug resistance problem [11]. While this strategy has already been reduced to practice in the case of pathogens resistant to β-lactams [40], efforts to develop inhibitors of resistance to aminoglycosides are still in experimental stages. We have recently found that ionophores complexed to Zn^+2^ or Cu^+2^ could be potentiators that decrease the levels of resistance to amikacin in *K. pneumoniae* and *A. baumannii* clinical isolates [23-25]. Since one of the ionophores that in complex with Zn^+2^ demonstrated activity as an inhibitor of the resistance to amikacin was CI8HQ, a substituted 8-hydroxyquinoline (8HQ), we expanded our studies to other compounds with these characteristics. Fig 1 shows the compounds tested in this work. The tests were carried out using as models three *A. baumannii* clinical isolates, two of them harboring the *aac(6’)-Ib* gene [32, 33]. The third strain, which does not carry this gene, exhibits resistance to amikacin by a different mechanism. Although this mechanism remains to be elucidated, it most probably consists of phosphorylation mediated by the *aphA6* gene found in its genome [33, 34].

**Fig 1.**
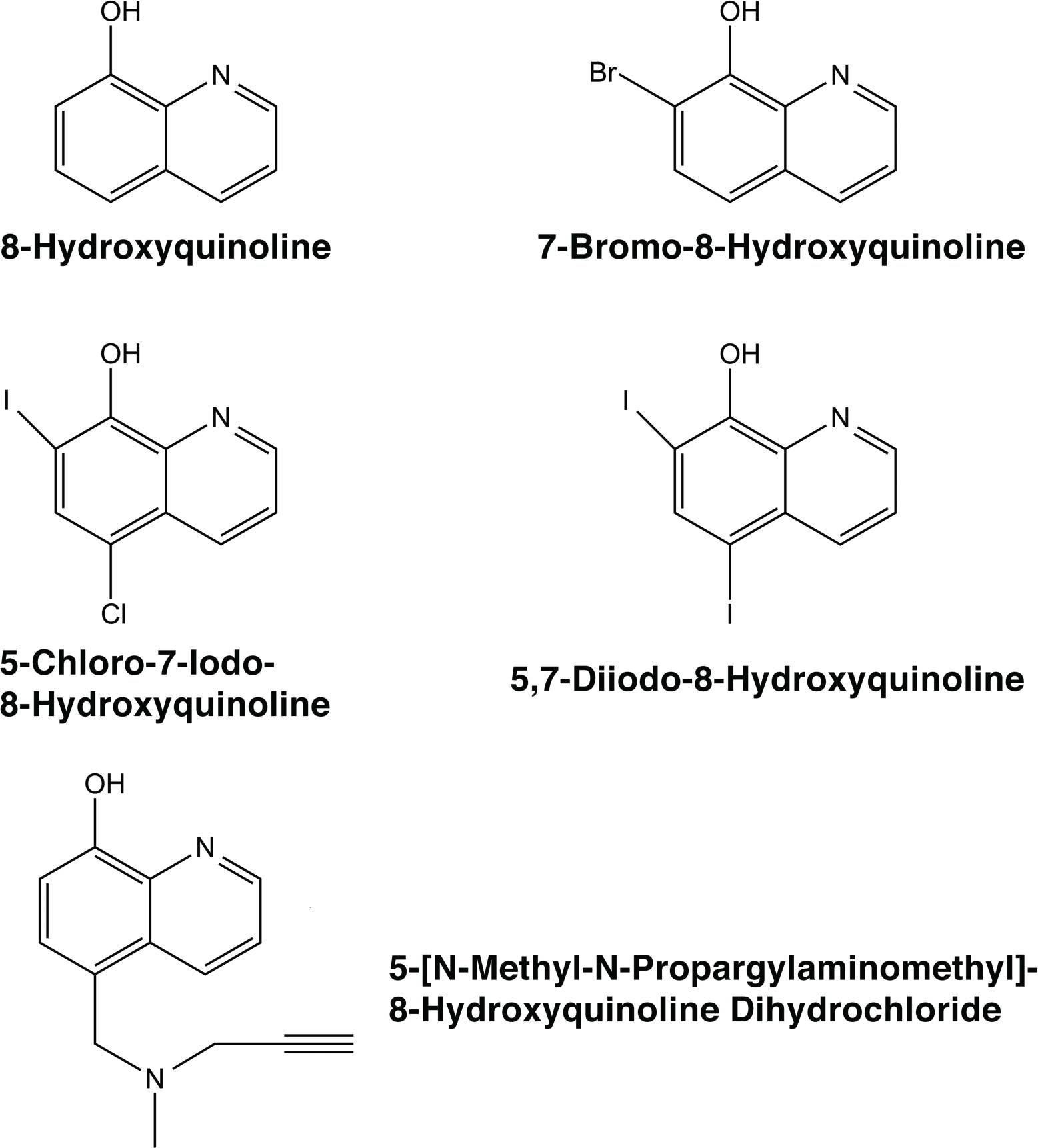
Chemical structures of 8-hydroxyquinoline and derivative compounds.

Growth curves in the presence of incremental concentrations of amikacin showed that the strains harboring *aac(6’)-Ib*, A144 and A155, can grow in up to 16 μg/ml of the antibiotic (S1 Fig, A and B). Conversely, strain Ab33405 had a different behavior, while the lag phase became longer as the amikacin concentration was increased, healthy growth was observed at all tested concentrations (S1 Fig, C). These results are in agreement with the finding that the latter strain resists amikacin using a mechanism different from that in strains A144 and A155. To confirm that *A. baumannii* Ab33405 is not able to mediate enzymatic acetylation of amikacin, the total soluble protein extracts of all three strains were used in *in vitro* acetylation assays using amikacin and AcetylCoA as substrates. Table 1 shows that while extracts from strains A144 and A155 mediated incorporation of radioactive acetyl groups to the acceptor substrate, the extract obtained from strain Ab33405 lacked acetylation activity.

**Table 1.**
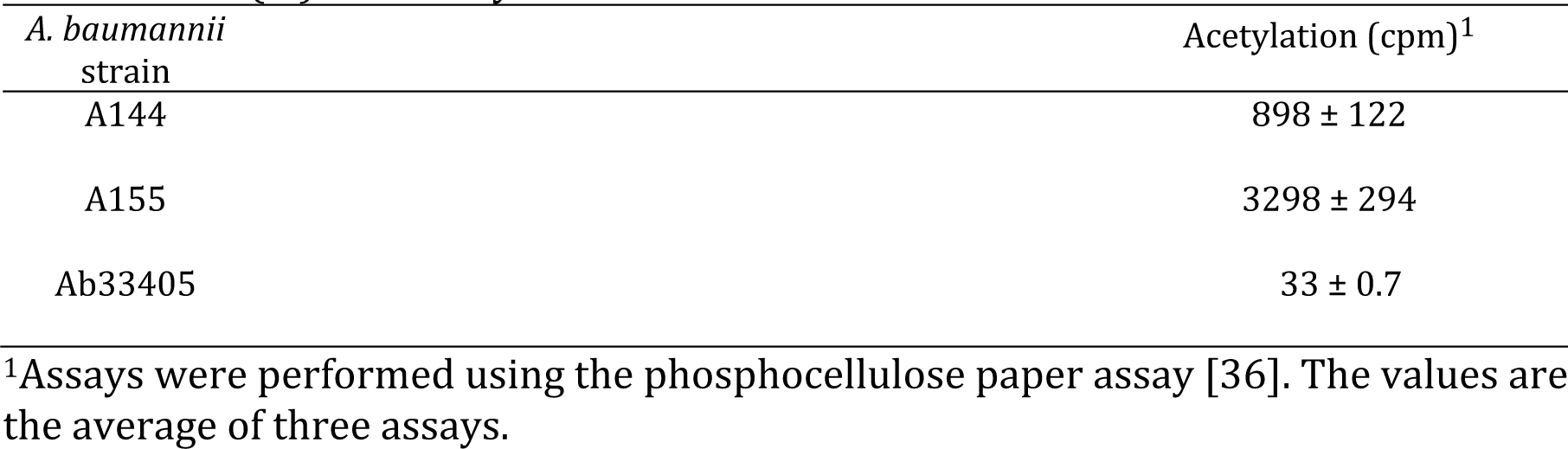
AAC(6’)-Ib activity

| *A. baumannii*<br>strain | Acetylation (cpm)<sup>1</sup> |
| --- | --- |
| A144 | 898 ± 122 |
| A155 | 3298 ± 294 |
| Ab33405 | 33 ± 0.7 |

<sup>1</sup>Assays were performed using the phosphocellulose paper assay [36]. The values are the average of three assays.

The growth of all three *A. baumannii* strains was unaffected by the presence of 25 or 50 μM ZnCl_2_ or up to 10 μM 8HQ, CI8HQ, 5-[N-Methyl-N-propargylaminomethyl]-8-hydroxyquinoline (MP8HQ), or 5,7-diiodo-8-hydroxyquinoline (II8HQ) (S1 Fig, A-C). Conversely, 10 μM 7-Bromo-8-hydroxyquinoline (B8HQ) was toxic to all three strains, and while strains A155 and Ab33405 could grow in the presence of up to 5 μM, strain A144 growth was inhibited at 1 μM B8HQ (S1 Fig, A-C).

Once concentrations of the ionophores and ZnCl_2_ that were not toxic to growing bacteria were identified, their activity as potentiators of amikacin was determined. These assays showed that CI8HQ and II8HQ were the only 8HQ derivatives that mediated phenotypic conversion to susceptibility to amikacin in strains A144 and A155 (Fig 2). Inspection of these results also showed that after 16 h, strain A155 started to grow when the ionophore tested was II8HQ. We do not yet have a satisfactory explanation for this observation. The ionophores 8HQ and MP8HQ were unable to induce any modification in the growth of strains A144 and A155 in the presence of amikacin and ZnCl_2_ (Fig 2). The tests where the ionophore used was B8HQ showed a reduction in growth in the presence of combinations that included B8HQ but either amikacin or ZnCl_2_ were omitted suggesting that the toxic effect of B8HQ is playing a role in growth inhibition rather than interference with acetylation of amikacin (Fig 2). Strain Ab33405 showed healthy growth in the presence of either of the ionophores plus amikacin and ZnCl_2_ confirming that the inhibition by Zn^+2^ is specific for resistance mediated by the modifying enzyme. Only one condition showed modest inhibition of growth (see Fig 2, strain Ab33405, CI8HQ) but some reduction in growth is also observed in the absence of ZnCl_2_, which may indicate unspecific inhibition. These results taken together with previous studies, especially those by Li et al. where the authors show than Zn^+2^ inhibits several modifying enzymes, indicate that ionophores complexed to metal ions can be an excellent strategy to interfere with resistance to aminoglycosides. However, this option might be effective only in cases of resistance mediated by selected aminoglycoside modifying enzymes. Interestingly, a recent report described that the metal homeostasis-disrupting action of ionophore-zinc complexes potentiates several antibiotics to restore susceptibility in resistant Gram-positive bacteria [41].

**Fig 2.**
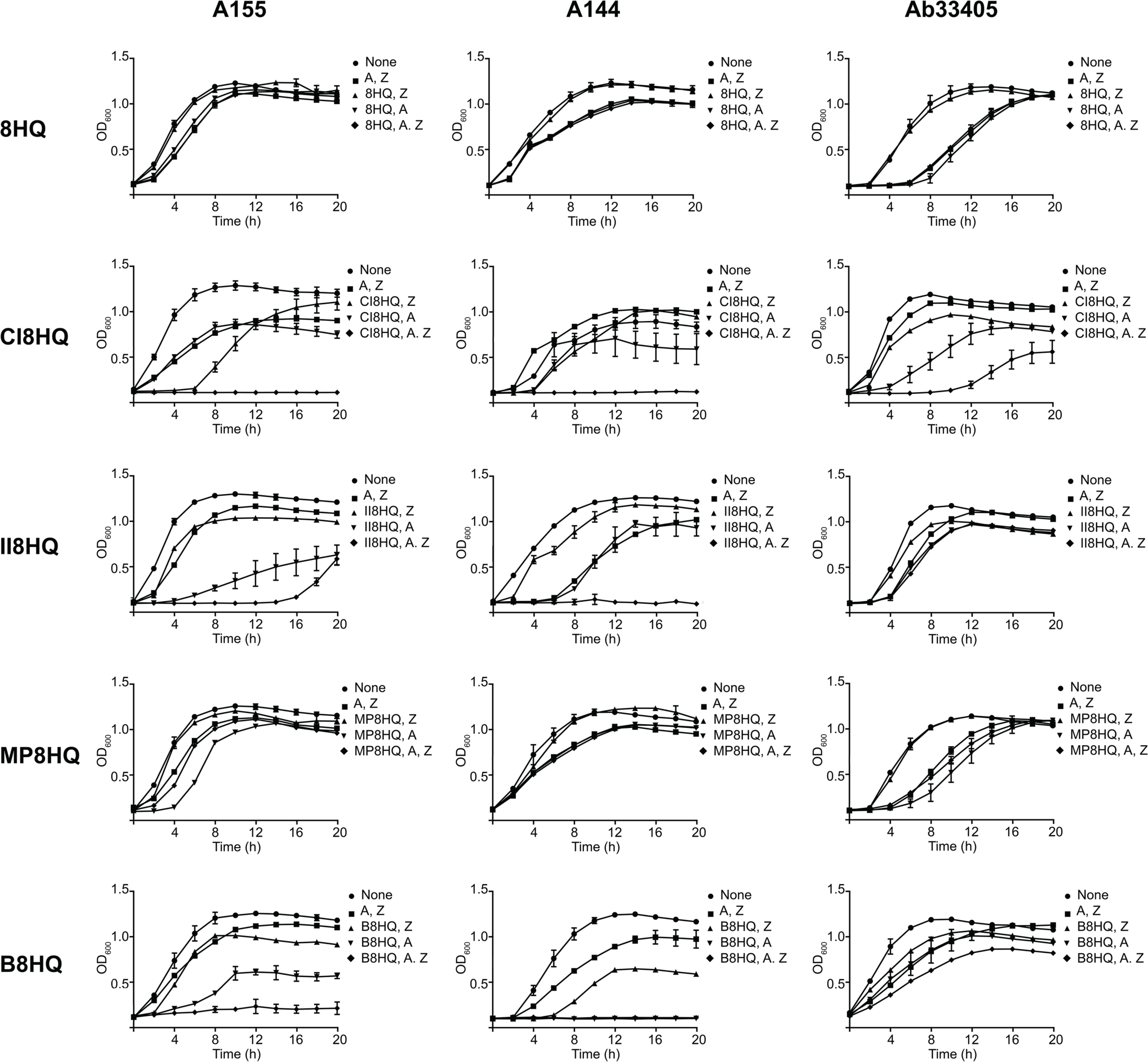
Effect of ionophore-zinc complexes on resistance to amikacin in *A. baumannii* strains. *A. baumannii* A155 (panels to the left), A144 (center panels) or Ab33405 (panels to the right) were cultured in 100 μl Mueller-Hinton broth in microtiter plates at 37°C, with the additions indicated in the figure and the OD_600_ was periodically determined. The concentrations used were 8 μg/ml amikacin, 25 μM ZnCl_2_, 5 μM ionophore. A, amikacin; Z, ZnCl_2_.

The results described above showed that CI8HQ and II8HQ were the most efficient ionophores that in complex with Zn^+2^ were able to mediate a conversion to susceptibility to amikacin in those *A. baumannii* strains in which resistance is mediated by AAC(6’)-Ib. The bactericidal effect of the combination ionophore-zinc and amikacin was confirmed using time-kill assays. Amikacin at a concentration as low as 8 μg/ml showed a robust bactericidal activity on *A. baumannii* A144 and A155 strains in the presence of the complexes (Fig 3). As expected, these strains did not lose viability when incubated with the antibiotic or any other combination of components that did not include all three of them (Fig 3). Also expected was the absence of bactericidal effect when the combinations ionophore-zinc plus amikacin were added to cultures of *A. baumannii* Ab33405 or the ionophore utilized was 8HQ (Fig 3). These results confirmed that amikacin can regain its bactericidal power in the presence of Zn^+2^ ions when resistance is due to AAC(6’)-Ib-mediated acetylation.

**Fig 3.**
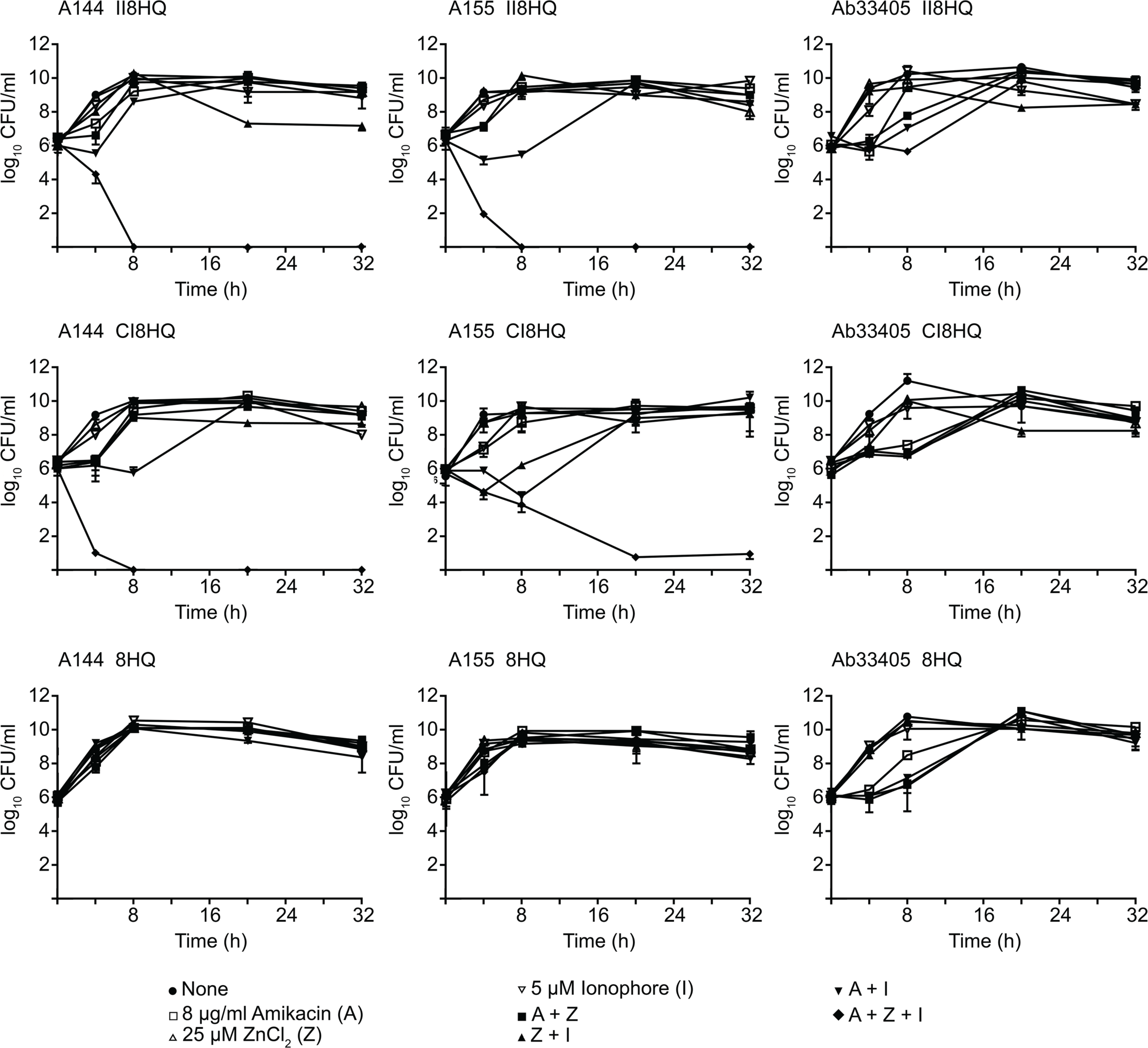
Time-kill assay curves for amikacin in the presence of ionophore-zinc complexes. *A. baumannii* A155 (panels to the left), A144 (center panels) or Ab33405 (panels to the right) were cultured in 100 μl Mueller-Hinton broth in microtiter plates at 37°C, with the additions indicated in the figure and the OD_600_ was periodically determined. A, amikacin; Z, ZnCl_2_; I, ionophore.

The ionophores tested in this work were used in a standard cytotoxicity assay as described in the Materials and Methods section. Addition of 8HQ, CI8HQ, or II8HQ, alone (S2 Fig) or in combination with amikacin and Zn_2_Cl to the cells did not result in significant toxicity (Fig 4).

**Fig. 4.**
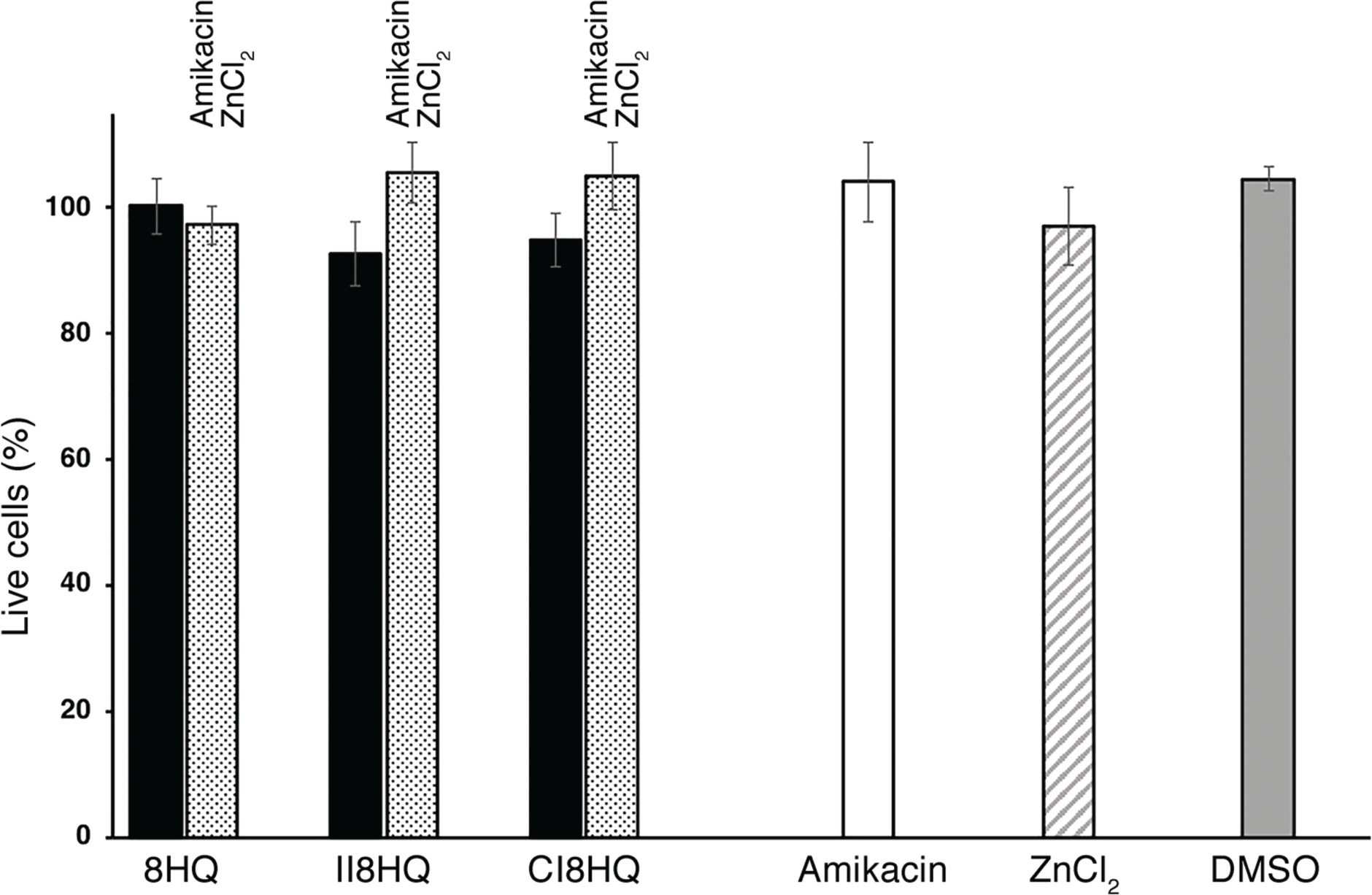
Cytotoxicity tests. Cytotoxicity on HEK293 cells treated with the indicated concentrations of the different compounds for 24 h was assayed using a LIVE/DEAD kit. The percentage of dead cells was calculated relative to the cells treated with DMSO. Cells incubated with 0.1% Triton X-100 for 10 min were used as a control for maximum toxicity. Experiments were conducted in triplicate and the values are mean ± SD. Black bars show survival in the presence of 5 μM ionophore. Stippled bars show survival in the presence of 5 μM ionophore, 25 μM ZnCl2, and 8 μg/ml amikacin. The same concentrations were used to determine survival in amikacin (white bar) and ZnCl2 (hatched bar). The concentration of DMSO used in the control was μM (gray bar).

## Discussion

Numerous approaches are being pursued to combat the current crisis of antibiotic resistance [11, 12]. In addition to the efforts to find or design new classes of antibiotics, researchers are looking for new scaffolds or attempting to modify existing antimicrobial families or designing compounds that act as adjuvant of these antibiotics by interfering with resistance [12, 42-46]. We have recently found that Zn^+2^, when complexed to ionophores such as pyrithione or CI8HQ, significantly reduces the levels of resistance to amikacin mediated by the AAC(6’)-Ib enzyme [23-25]. Since this enzyme is the most prevalent in amikacin resistant infections in the clinics [6], this finding represented a significance advance in the search for compounds that in combination with the antibiotic can help extend its useful life. The obvious possibilities of these compounds as part of formulations composed of amikacin and the inhibitor warrant further research to find the best ionophores. Since CI8HQ is a derivative of 8HQ, in this work we tested combinations of Zn^+2^ with 8HQ and other commercially available derivatives. While CI8HQ and II8HQ show similar capacity to reverse resistance to amikacin, 8HQ and MP8HQ did not show any of the desired inhibitory activity, and B8HQ exhibited antimicrobial activity in the absence of the antibiotic. The disparity of effects found among these chemically related compounds shows the importance of assessing the activity of ionophores with similar structures. Since one of the most crucial problems exhibited by numerous compounds that are otherwise good drug or adjuvant candidates is their toxicity, it was interesting that the ionophores tested in this work did not show cytotoxicity in our assays. Furthermore, as they are being researched as treatments of other human conditions, their low toxicity has also been established by other laboratories. Taken together, the results described in this work indicate that Zn^+2^ or other cations, complexed to ionophores are firm candidates to be developed as potentiators to aminoglycosides to overcome resistance, in particular CI8HQ and II8HQ are excellent candidates as adjuvants to overcome AAC(6’)-Ib -mediated resistance to amikacin.

## Supporting information

porting information Fig S1 and S2

## Acknowledgements

This work was supported by Public Health Service grant 2R15AI047115 from the National Institute of Allergy and Infectious Diseases, National Institutes of Health.

## Supporting information

**Fig S1. Effect of addition of different reagents on growth of *A. baumannii* strains.**

**Fig S2. Cytotoxicity tests.**

