## Supplementary material for "Restoration of susceptibility to amikacin by 8-hydroxyquinoline analogs complexed to zinc": porting information Fig S1 and S2

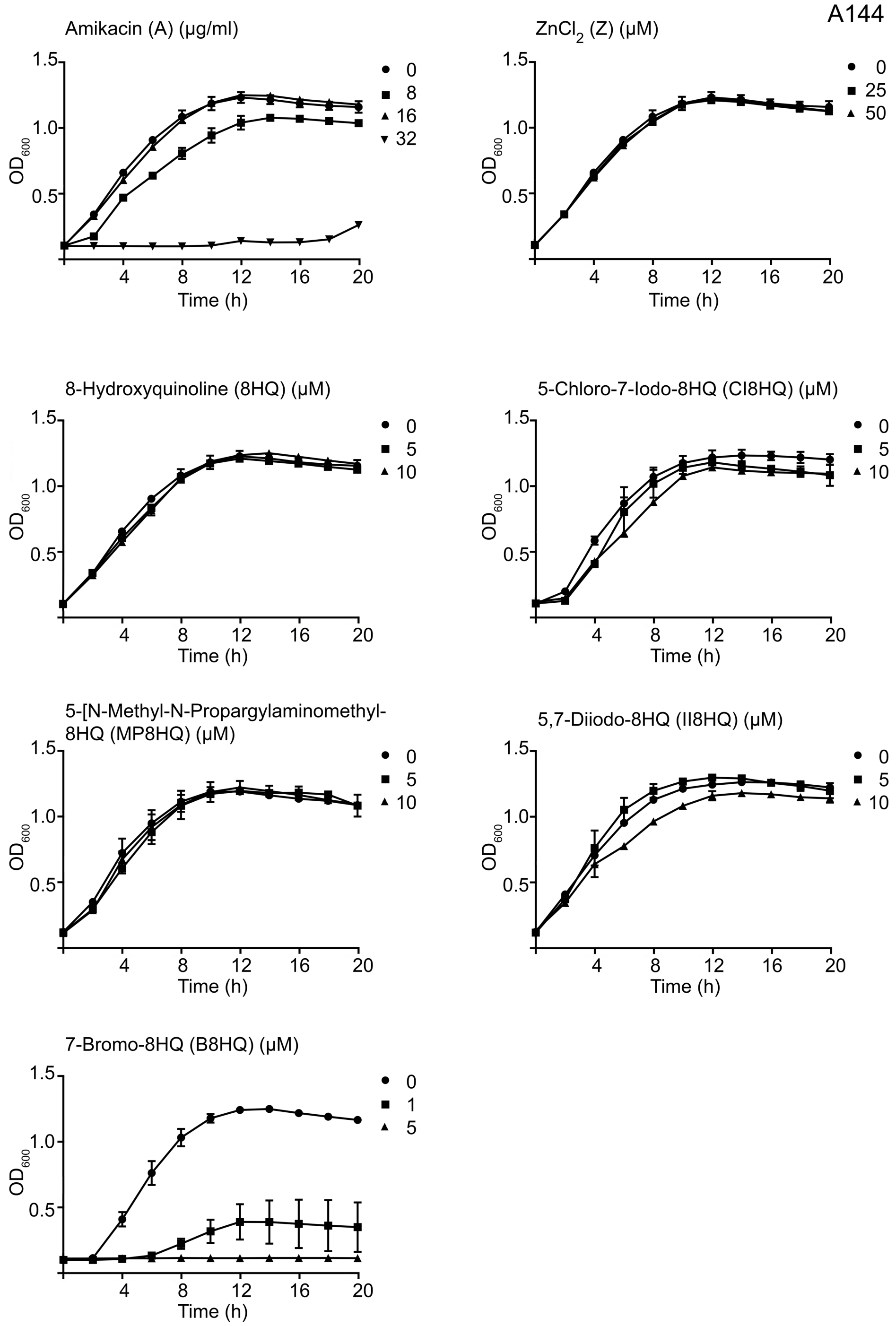
A144
Amikacin (A) (μg/ml)
0
8
16
32
ZnCl2 (Z) (μM)
0
25
50
8-Hydroxyquinoline (8HQ) (μM)
0
5
10
5-Chloro-7-Iodo-8HQ (CI8HQ) (μM)
0
5
10
5-[N-Methyl-N-Propargylaminomethyl-8HQ (MP8HQ) (μM)
0
5
10
5,7-Diiodo-8HQ (II8HQ) (μM)
0
5
10
7-Bromo-8HQ (B8HQ) (μM)
0
1
5
OD600
1.5
1.0
0.5
Time (h)
4
8
12
16
20

S1 Fig, B

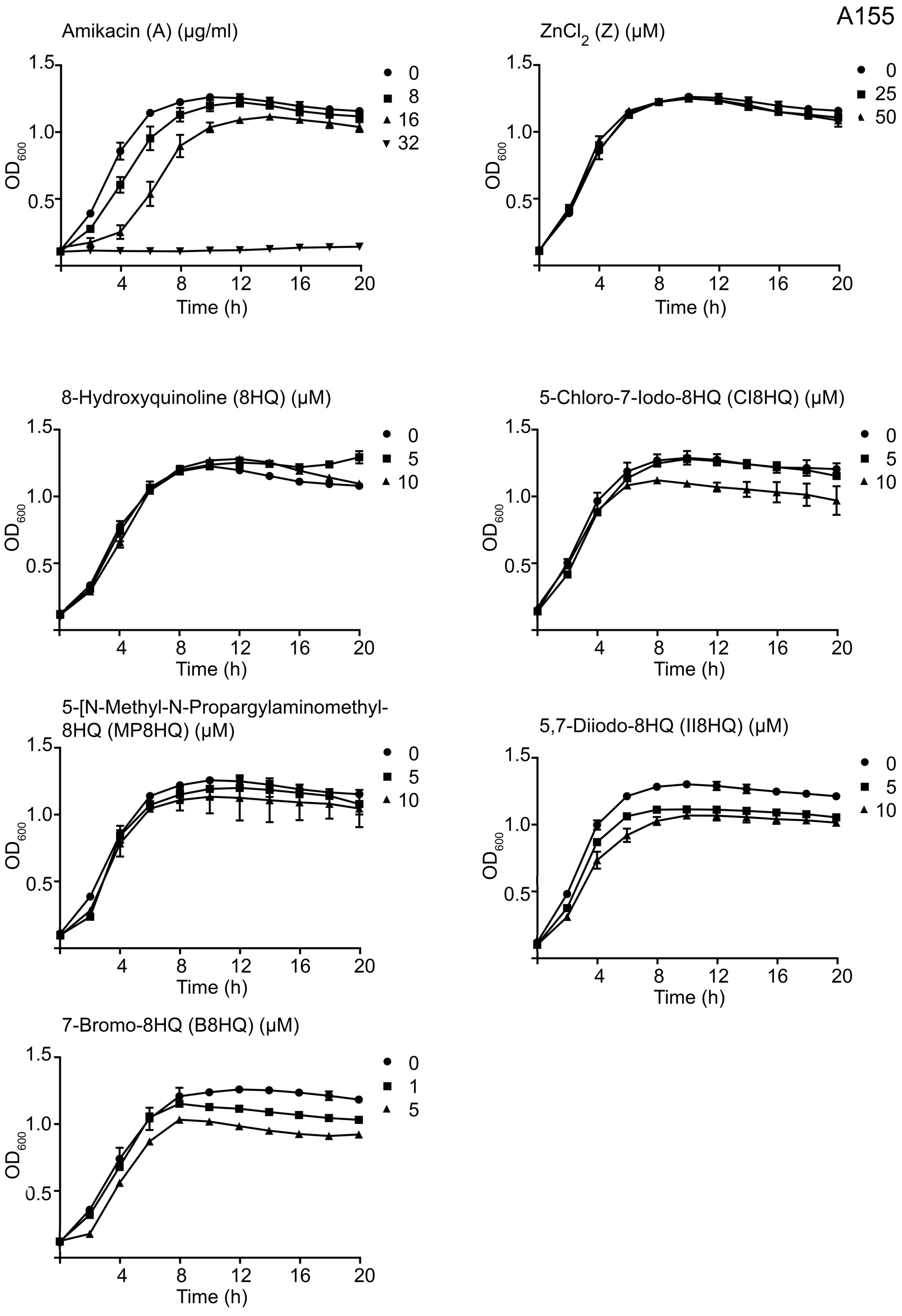
A155
Amikacin (A) (μg/ml)
0
8
16
32
ZnCl2 (Z) (μM)
0
25
50
8-Hydroxyquinoline (8HQ) (μM)
0
5
10
5-Chloro-7-Iodo-8HQ (CI8HQ) (μM)
0
5
10
5-[N-Methyl-N-Propargylaminomethyl-
8HQ (MP8HQ) (μM)
0
5
10
5,7-Diiodo-8HQ (II8HQ) (μM)
0
5
10
7-Bromo-8HQ (B8HQ) (μM)
0
1
5
OD600
Time (h)
0.5
1.0
1.5
4
8
12
16
20

S1 Fig, C

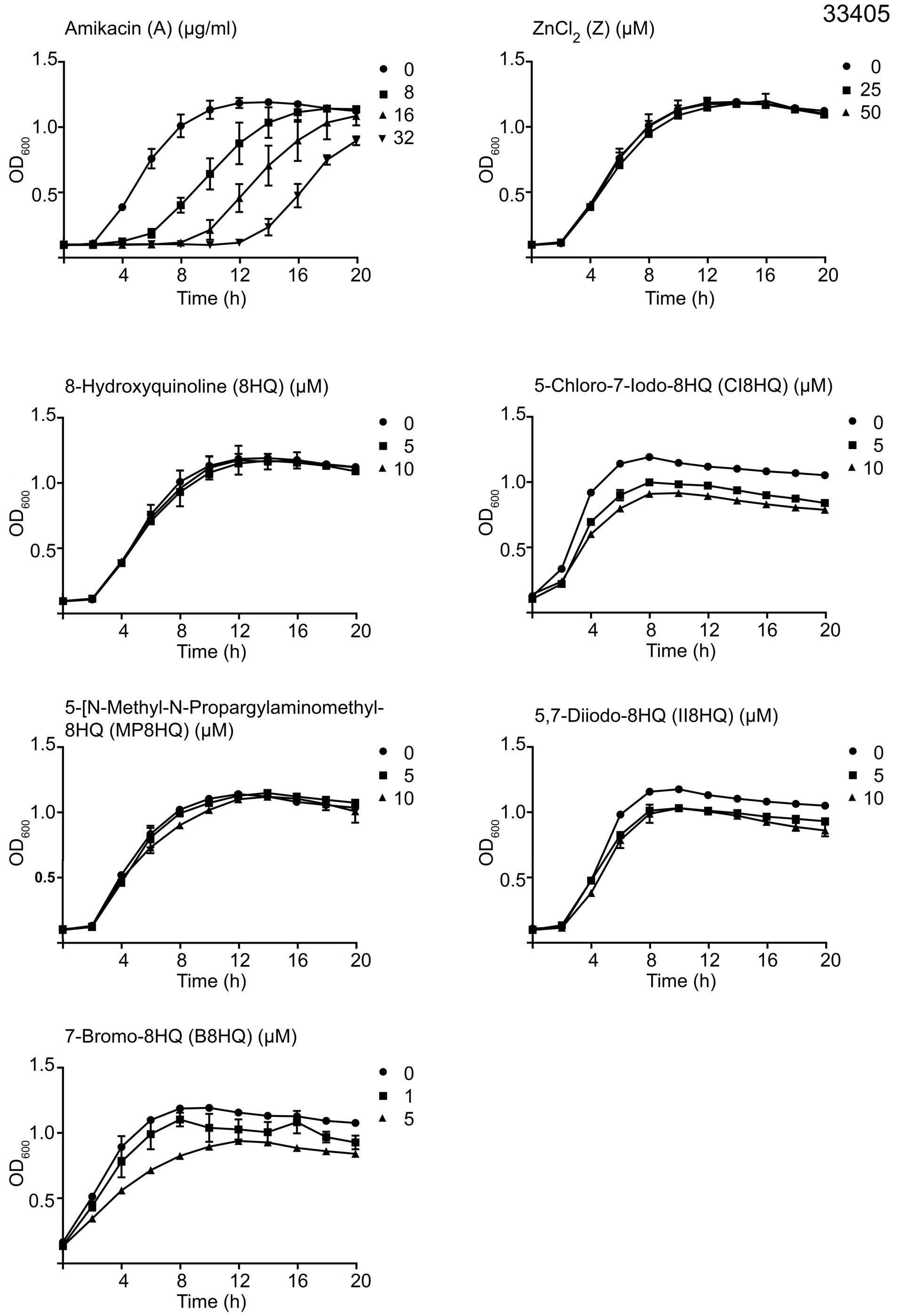


**Fig S1. Effect of addition of different reagents on growth of *A. baumannii* strains.** *A. baumannii* A144 (A), A155 (B) or 33405 (C) were cultured in 100 µl Mueller-Hinton broth in microtiter plates at 37°C, with the additions indicated in the figure and the $OD_{600}$ values were periodically determined.

S2 Fig

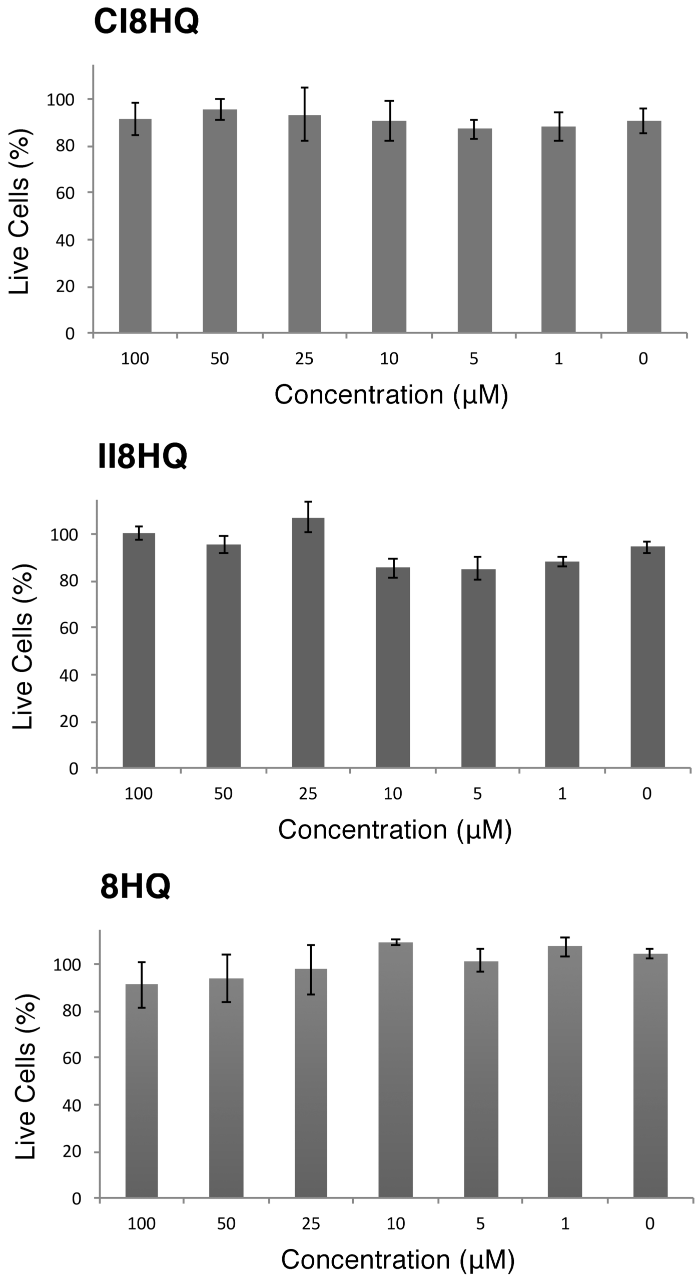


**Fig S2. Cytotoxicity tests.** Cytotoxicity on HEK293 cells treated with the indicated concentrations of the different compounds for 24 h was assayed using a LIVE/DEAD kit. The percentage of dead cells was calculated relative to the cells treated with DMSO. Cells incubated with 0.1% Triton X-100 for 10 min were used as a control for maximum toxicity. Experiments were conducted in triplicate and the values are mean ± SD (n=5).
